# Deep Archetypal Analysis for interpretable multi-omic data integration based on biological principles

**DOI:** 10.1101/2024.04.05.588238

**Authors:** Salvatore Milite, Giulio Caravagna, Andrea Sottoriva

## Abstract

High-throughput multi-omic molecular profiling allows probing biological systems at unprecedented resolution. However, the integration and interpretation of high-dimensional, sparse, and noisy multimodal datasets remains challenging. Deriving new biology using current methods is particularly difficult because they are not based on biological principles, but instead focus exclusively on a dimensionality reduction task. Here we introduce MIDAA (Multiomic Integration with Deep Archetypal Analysis), a framework that combines archetypal analysis, an approach grounded in biological principles, with deep learning. Using the concept of archetypes that are based on evolutionary trade-offs and Pareto optimality – MIDAA finds extreme data points that define the geometry of the latent space, preserving the complexity of biological interactions while retaining an interpretable output. We demonstrate that indeed these extreme points represent cellular programmes reflecting the underlying biology. We show on real and simulated multi-omics data how MIDAA outperforms state-of-the-art methods in identifying parsimonious, interpretable, and biologically relevant patterns.

## Main text

Fundamental processes in cellular biology such as cell differentiation, development, and carcinogenesis are inherently driven by multiple interacting molecular layers. Those encode the information that orchestrates the intricate regulatory networks of proteins, transcription factors, and signalling molecules^2^ that give rise to biological phenomena Any attempt to look at a single molecular layer at a time is bound to miss crucial biological insights. High-throughput multi-omics technologies that can measure many concurrent molecular layers in the same cell or sample promise to help gaining a more comprehensive picture of biological phenomena^3^. However, integrating and extracting patterns from these types of high-dimensional, noisy, and sparse data is a major statistical and algorithmic challenge^4^. The biggest problem is that current state-of-the-art methods are not based on biological principles, but merely focus on the problem of dimensionality reduction in a data-driven fashion. This makes the output of those approaches hard to interpret from a biological perspective, and they do not allow inferring the putative causes of observed changes in biology.

For example, one of the key steps in integrating multiple data sources is dimensionality reduction, with algorithms specifically developed to integrate biological data^5^. For instance, probabilistic multi-omics factor analysis (MOFA) has been successfully applied to find relevant patterns from multiple omics data sources (e.g., gene expression, protein abundances)^6–8^. However, MOFA and other factor analysis methods posit that the observed data can be linearly reconstructed from the latent factors and their corresponding loadings. Linear models simplify the mathematical treatment and their latent factors are interpretable in relation to the measured data. For instance, a factor with high loadings from genes involved in a specific metabolic pathway might be interpreted as representing that pathway. However, linear models miss complex non-linear interactions typical of real biological systems, such as the non-proportional relationship between gene expression and metabolite concentrations^9^, threshold-dependent effects of epigenetic modifications on gene activity^10^, cooperative transcription factor binding^11^, and general environmental factors^12^.

In computational biology, a popular non-linear dimensionality reduction framework is the Variational Autoencoder (VAE) architecture^13,14^, which can model arbitrarily complex non-linear interactions between the input variables via an encoding/decoding mapping parametrized by a deep neural network. The latent space provided by VAEs is more powerful and expressive than linear ones, yet it is no longer easy to interpret, making these models practically behave like “black-box” information compression machines^15^. This is a limitation for biological applications where we want to understand the system. In biology, we need generative models with an interpretable latent space that can be used to analyse specific system perturbations. Importantly, we need to inject biological principles into data integration approaches.

### MIDAA

Archetypal Analysis (AA)^16^ is a matrix factorization algorithm designed to decompose the input data as a convex (i.e., linear) combination of extreme data points called archetypes. Contrasted with other methods, AA forces strong constraints on the geometry of the latent space and recovers a set of bases that are expressed only in terms of the relative distances from the archetypes. Archetypal Analysis is grounded in the biological principles of evolutionary trade-offs and Pareto optimality, where extreme geometrical points in the space of biological ‘states’ represent phenotypic programmes cells or organisms converge to. AA is a promising alternative for dimensionality reduction because, by construction, its coordinate system is trivially interpretable in the very same domain of the data. The linear latent space of AA can be turned into a non-linear manifold by combining AAs with deep neural networks for archetypal decomposition^17^. In this way, it is possible to retain the interpretability of the latent space while leveraging the power of a non-linear dimensionality reduction. Building on this idea we developed MIDAA (Multiomic Integration with Deep Archetypal Analysis), an open-source

Python framework for the integration of multi-omics data with deep archetypal analysis. MIDAA supports different input types and neural network architectures, adapting seamlessly to the high complexity of modern biological data, which ranges from counts in sequencing assays to binary values in CpG methylation assays. In principle, the model could be extended to combine data from non-omics sources (text and images) when combined with embeddings from other deep-learning models. MIDAA is implemented on a PyTorch^18^ backend that leverages GPU acceleration, scaling to thousands of cells (e.g. 100,000 cells in ∼5 minutes for 500 epochs, Supplementary Figure 1).

**Figure 1.**
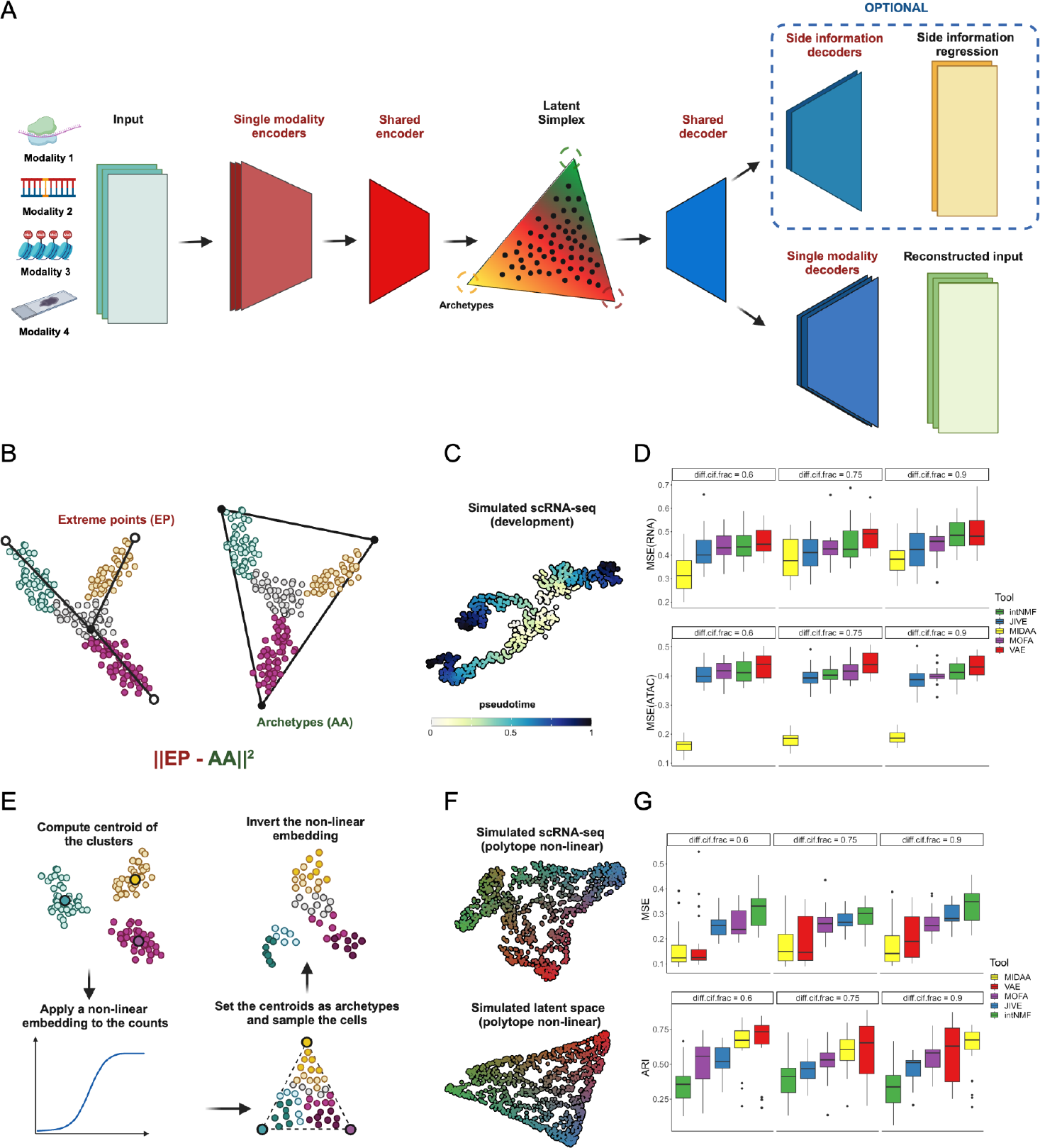
Performance of MIDAA on a multiomics benchmark dataset. **A**. Schematic representation of the model. We allow an arbitrary number of modalities in input, the model then encodes each modality using a private encoder. The last layer of these modality-specific encoders is concatenated and given as input to a shared encoder that learns the latent space and the simplex structure. The decoding part is exactly the reverse with the addition of an optional decoding branch for regression/prediction tasks. **B-C**. We simulate a branching differentiation process. We model the differentiation starting from a stem center population with pseudotime 0 differentiating towards 3 different states. Our goal here is to understand if the terminal (i.e. low and high pseudotime) state of differentiation is recapitulated correctly by the archetypes **D**. We measured the Mean Squared Error (MSE) between the aggregated expression (top panel) or peak counts (bottom panel) of cells at terminal states (bottom 15% and top 75% percentile of pseudotime) and the reconstructed archetypes **E-F**. For the second test, we sample from a simplex structure in a non-linear latent space, the non-linearity is parametrized by a neural network. **G**. Here we measure how well the tools reconstruct the original latent space. As error measures, we computed the MSE of the true and inferred archetype weights (top) and the Adjusted Rand Index (ARI) for the true and inferred highest archetype assignments. In all the plots “diff.cif.fraction” controls the fraction of divergence among archetypes or populations in the development trajectory, a lower number implicates a higher noise.

### Results on simulated data

Using synthetic data (Methods) we tested if MIDAA could decipher two relevant biological processes: cellular differentiation (Figure 1B,C) and evolutionary dynamics on a fitness landscape (Figure 1E,F). In both cases, we generated synthetic data from 1,000 cells (20 datasets, 3 noise levels) with matched ATAC/RNA sequencing (chromatin accessibility and gene expression measurements) and compared MIDAA to a pipeline where we first performed dimensionality reduction with both linear and non-linear models, followed by canonical AA. In the differentiation test, MIDAA largely outperformed competing methods, on average reducing the RNA and ATAC reconstruction error by 15% and 55%. In the evolutionary dynamics test dataset, MIDAA decreased the reconstructor error for the latent space by 13% (average), across all noise levels (Figure 1D-H). Notably, in the latter test, a clear performance difference was observed between linear and non-linear statistical models (Figure 1H), with MIDAA being the top performer on average. Interestingly, in the oversimplified case of a linear generative latent space (Supplementary Figure 2), while linear models achieved the lowest reconstruction error, MIDAA was the best non-linear model, suggesting its geometrical constraints regularise the model.

**Figure 2.**
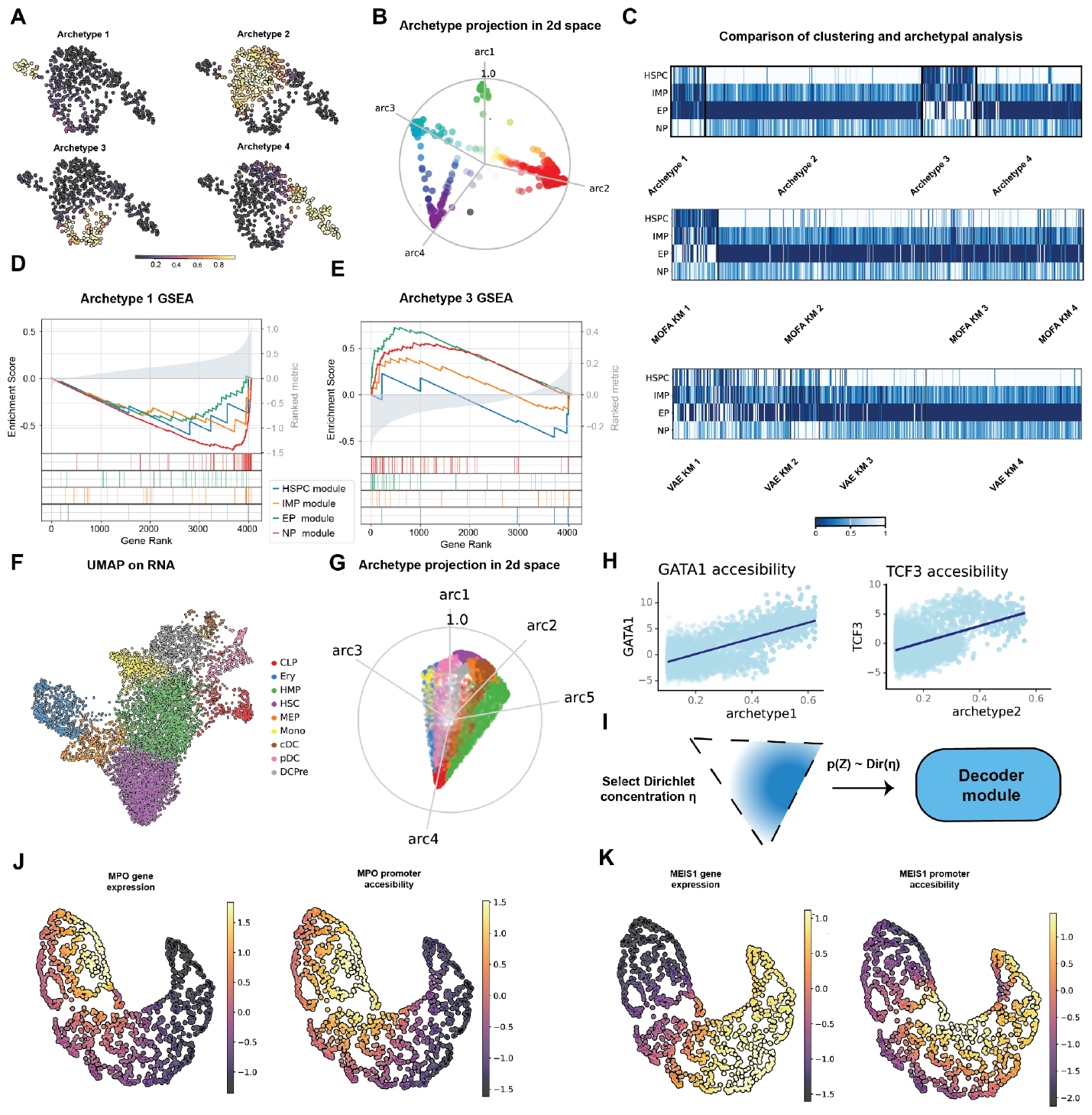
MultimodalDeep Archetypal Analysis reconstructs an efficient and biologically meaningful latent space. **A**. Archetype distribution plotted over the RNA UMAP **B**. A 2d projection of the simplex latent space. **C**. Heatmap of normalized [0-1] cell progenitor scores for cells with archetype probability >=80% and K-means clustering in VAE and MOFA space. **D-E**. GSEA enrichment analysis for archetypes 1 and 3 using the cell progenitor gene sets from ^19^. **F-G**. UMAP and 2d simplex projection of the dataset in ^20^. **H**. Correlation of transcription factor motif deviation and archetype weights. GATA 1 is an erythropoietic commitment marker and TCF3 is enriched in dendritic progenitors **I**. The generative nature of the model makes it easy to produce synthetic datasets from the latent space. First of all the user can sample from a Dirichlet distribution specifying the concentration parameter and from that the decoder generates realistic multi-modal data. **J-K**. Concordance of gene expression and promoter accessibility in a synthetic dataset consisting mainly of the erythropoietic and stem archetypes.

### Results on real data

A key problem that involves complex multimodal interactions is the differentiation of hematopoietic stem cells (HSC) into mature blood cells, known as hematopoiesis. We used MIDAA to extract biologically interpretable insights from single-cell multi-omics data (whole genome CpG methylation status and transcriptional activity) of CD34+ positive cells, a type of hematopoietic progenitor cell^19^. First, we calculated the level of commitment for specific lineages in each cell by computing a score for the gene signature presented in^19^. In particular, in this dataset we have a group of Hematopoietic Stem and Progenitor Cells (HSPC) differentiating first into Immature Myeloid Progenitors (IMP) and then into Erythroid Progenitors (EP) and Neutrophil Progenitors (NP). MIDAA did find 4 optimal archetypes from 512 cells (Supplementary Figure 3 and Figure 2A,B), and its latent space did recapitulate lineage commitment in this dataset. In particular, the archetype weights were found to be strongly associated with all the terminal states in the gene signature (Supplementary Figure 4), suggesting that the latent space geometry matches the one of the differentiation landscape. Notably, MOFA and VAE embeddings failed to extract the patterns of EP and NP cells, with the most relevant MOFA factors driven by highly variable samples. Overall, none of the competing methods managed to fully recapitulate the differentiation features of these cells (Supplementary Figure 5-6).

We then asked whether our latent space reproduced known differentiation lineages for these cells. To answer this, we compared our archetypes (recapitulated by cells with weight >80%) to a k-means clustering in MOFA and VAE latent spaces. Our archetypes identified clear progenitor cells, whereas the standard achieved the worst separation (MIDAA silhouette score increased by ∼90%) (Figure 2 C and Supplementary 7). Interestingly, this analysis highlighted that Immature Myeloid Progenitors (IMP) were not represented by a single MIDAA archetype, but rather by a combination of them. This is consistent with IMP cells being in a transition state from Hematopoietic Stem and Progenitor Cells (HSPC) to Erythroid Progenitors (EP) and Neutrophil Progenitors (NP).

Finally, we tested whether archetypes could be used in an unsupervised way for the discovery of biological programs. To investigate this, we ran a gene set enrichment analysis on the expression reconstructed for each archetype, using as input gene sets the 4 cellular programs of hematopoietic progenitors. We found one archetype enriched for genes characteristic of EP, and one positively enriched for NP genes while negatively enriched for HSPC genes (Figure 2D,E), consistent with our previous clustering analysis. This suggests that MIDAA’s archetypes can be easily associated with well-defined biological characteristics, and that can be used for downstream analysis as representative of real data measurements.

To further confirm MIDAA’s flexibility and potential to adapt to different input data types, we analysed a different cohort of CD34+ cells, this time generated with 10x GEX + ATAC libraries^20^. MIDAA found an optimal number of 5 archetypes. (Figure 2F,G). Taking advantage of the ATAC measurement, we first ran chromVar^21^ to calculate transcription factor (TF) motif deviations in the dataset. We then correlated the inferred values for some key TFs of hematopoietic development with the archetype weights, observing a significant positive correlation trend (Figure 2H). This shows again how MIDAA’s latent space recapitulates the known biological processes in the data.

### Simulating realistic synthetic multi-omics datasets

Thanks to its generative architecture, MIDAA makes it possible to simulate multi-omics data from the latent space in a biologically informed way. The user is free to sample from the latent simplex by weighing the importance of the different archetypes. From these samples, the decoder will then generate realistic observable data (Figure 2I). To show this, we sampled a synthetic dataset consisting mainly of archetypes 3 and 5, associated respectively with HSC and dendritic cell (DC) progenitors. The output recapitulates exactly the expected HSC to DC transition, as evidenced by the MPO and MEIS1 markers. Notably, this effect is observed both at the gene expression level and as chromatin accessibility in the promoter, proving how MIDAA can produce realistic synthetic data that are consistent across modalities.

## Conclusions

Collectively, these findings demonstrate that MIDAA generates interpretable, biologically coherent, and expressive embeddings for multi-omics data. Moreover, thanks to its generative architecture MIDAA can also be used to simulate new synthetic data. MIDAA architecture is naturally modular and easily generalizable and in the future, we envision a broad applicability of this methodology also to other types of relevant multi-modal data, such as images as well as protein and DNA sequence embeddings.

## Methods

### The Matrix Factorization Problem

Omics data is commonly represented in the form of high dimensional, sometimes sparse, numerical matrices. In this context, dimensionality reduction becomes not only essential to make subsequent analysis feasible from a computational point of view, but also to filter out technical noise and minor sources of variability. Indeed, the most common analysis pipeline for scRNA-seq and ATAC-seq data involves first a dimensionality reduction step using PCA or LSI and then graph modularity clustering to extract relevant groups in the dataset.

The general definition of the problem is quite simple, given an input matrix **X**^**NxM**^ where *N* ∈ ℕ represents the number of the samples and *M* ∈ ℕ the number of features, we wish to find a two matrix decomposition of **X**. In other words, after fixing an *R* ∈ ℕ < *max*(*N,M*), our decomposition writes as:

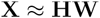

Where **H** is an *N* x *R* matrix and **W** is an *R* x *M* matrix. This formulation describes an extremely broad family of methods, the specific constraints and properties we force on the two matrices **H** and **W** as well as the metrics we want to optimize for the reconstruction.

In case we have multiple input modalities, if we index them by *g* = [1,…,*G*] we can naturally reframe the problem as:

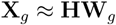

In this case we allow the number of features to differ by modality so that we have **X**_*g*_ and **W**_***g***_ specific for each modality with dimension *N* × *M*_*g*_ and *R* × *M*_*g*_ where *M*_*g*_ is the number of features for modality *g*.

### Archetypal Analysis

Archetypal Analysis (AA), is a dimensionality reduction method that solves the matrix factorization problem by enclosing the data into a convex polytope. The vertices of this polytope which spans the convex hull of the polytope are called archetypes and are generally interpreted as extreme or ideal samples in the dataset.

More formally, let us fix the number of vertices (or equivalently archetypes) to. *K* ∈ N We then introduce the matrices **A** = (*a*_*nk*_) and **B** = (*b*_*nk*_) with sizes respectively of *N* × *K* and *K* × *N*. Moreover we constraint these matrices to be row stochastic, namely:

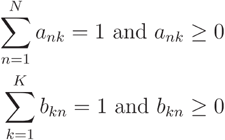

Finally, our AA decomposition reads, by assuming again multiple input modalities:

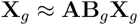

Note that if we set *R* = *K*, **H** = **A** and W = B_*g*_X_*g*_ we recover the original matrix factorization problem.

The original algorithm to solve this was introduced by ^16^ and was formulated as an alternating least square problem on the two matrices **A** and **B**. Faster approaches have been developed such as the Principal Convex Hull method ^22^ and the Frank-Wolfe method ^23^ which are conceptually based on gradient descent. Nevertheless, also those former optimized methods while faster, still scale proportionally to the full input size, becoming slow for big matrices. Archetypal analysis has been successfully used in modeling single-modality data in biology^23–25^, our goal here is to extend it to multimodal data and provide a unified framework in the context of deep latent variable models. All of this is conveniently packed in a user-friendly Python package that easily adapts to the plethora of omics data currently available.

### Deep Multiomics Archetypal Analysis

We started from the Deep Learning extension of the Archetypal Analysis proposed in ^17^ to build our MIDAA model. Our main goal is to perform amortized inference over the two matrices **A** and **B** in some latent space **Z**. Ideally, we want to have a reduced latent representation of the input in some non-linear shared space **Z** and then learn the convex polytope. Indeed, our method performs joint inference over the polytope and the latent space. To reduce the degrees of freedom and avoid optimising both over the number of archetypes and the dimensionality of the hidden space, we fix the polytope shape to be a simplex and set the number of dimensions of the hidden space as the number of archetypes - 1, as in ref (cite deep AA). We will use an encoder-decoder to encode our latent space and project back the AA results.

Formally, let us define the number of latent dimensions as K - 1 and the latent space representation as **Z** with dimensions *K* × *N*, then we can define the simplex reconstruction in latent space **Z**^**\***^ as:

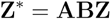

Differently from ^17^ we do not fix **BZ** to be the standard simplex but we explicitly learn and compute both **B** and **Z** in one passage.

We do this both to simplify the model as we have less parameters to tweak and because our latent space formulation achieves better average scores on our synthetic tests (Supplementary Figure 9). In our model, we constrain **Z** to be in [0,1]^*k*-1^ instead of the standard isotrophic gaussian used in VAEs ^26^. This choice will be made clearer in the section regarding inference.

We employ an encoder-decoder inference strategy, so the 3 matrices are the output of a function 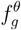 parametrized by a neural network, the same for the decoder. To make this paragraph more readable we keep a single function for the encoder. Still, it is worth noticing that the first encoding part is modality specific (i.e. an independent function and network). Then the output of each single modality encoder is concatenated and given in input to a shared one, as illustrated in Figure 1A.

To compute the loss we project back the simplex reconstruction in the original space using the decoder. We have one principal part of the loss which is the usual input reconstruction; in addition, we allow the network to classify side data **Y**, that we index with *s* ∈ ℕ. This is useful when we want our archetypes also to reflect some external variables that we might not want to include in the encoding phase.

In particular, given a likelihood distribution with its parameter set, the total likelihood reads as:

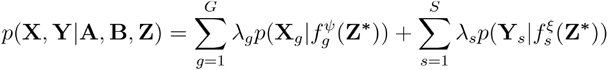

Where we define **X** = [**X**_**1**_,…,**X**_**G**_] and **Y** = [**Y**_**1**_,…,**Y**_**G**_]. Here 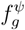 and *f*_*s*_ are the decoding network for the side and input data. Again for simplicity, we omit that there is a shared part and a modality-specific part and refer to Figure 1A. As different modalities can have different numbers of features we allow the user to specify constants *λ*_*g*_ and *λ*_*s*_ to normalize the likelihood, by default, they are set to respectively 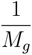 and 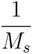 give the same importance to each modality (where *M*_*g*_ and *M*_*s*_ are the number of features for each input and side modality).

### Model Inference and formulation

We define the learning objective in a way akin to that of a standard VAE^26^ but with some major differences regarding the form of the distribution involved in the latent space. The loss function however that we optimize is the same and it is the evidence lower bound (ELBO) that we maximize throughout training using stochastic gradient descent with the Adam optimizer:

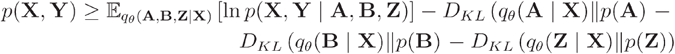

Our variational distributions q is defined over the matrices **A, Z**, and **B** and we assume a mean-field factorization as 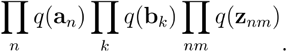. To keep the notation consistent for **A** and **B** here we multiply over respectively the rows, the columns (i.e. we assume independence among archetypes, latent dimensions, and samples).

The choice of the variational distributions comes naturally from the constraint of AA:

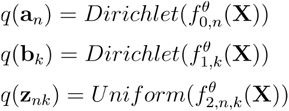

Here with 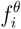 we index the output dimensions of the encoder. Priors have the same functional forms as the variational posteriors and have equal unitary concentration for the Dirichlet while the Uniform has range [-1,1]. Regarding the distribution of **Z** we departed from the standard isotropic Gaussian as a prior as it tends to concentrate probability density on the shell of a hypersphere (in high dimension) or push towards the center (in lower dimensions) and as such makes the space particularly bad suited for learning a simplex representation of the data^27^.

Regarding the likelihood distributions we allow flexibility and currently support the following distributions as valid likelihoods: **Beta, Poisson, Gaussian, Gamma, Negative Binomial**, and **Categorical**.

The integration of new likelihood is easy given the modularity of the model and we plan on increasing the distribution support following requests from the community. In all cases, parameters are amortized by the network 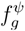 for input reconstruction and the 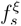 for side data.

### Benchmark on simulated data

We used the scMultisim tool^28^ to generate synthetic single-cell multi-omics data, which uses real-data inferred gene regulation networks to sample both trajectory-like and clustered gene expression and chromatin accessibility data.

As a comparison we choose a set of popular methods in multi-omics data integration, spanning a wide range of different statistical techniques: JIVE^29^ based on PCA, intNMF^30^ based on NMF, MOFA^7^ based on Factor Analysis, and a vanilla VAE^26^. In the latent space produced by each of these methods, we then ran linear archetypal analysis as implemented in the R package ^31^.

We analyzed two main case studies: one in which the latent space is a simplex and one in which it is instead a differentiation trajectory. In the first case, we generated a cohort in which the mapping function from the space of observables to the latent space is linear and another one in which it is non-linear.

For each of these cases, we simulated 20 datasets of 1000 cells.

We also repeated the experiments for three values of the parameter *diff*.*cif*.*fraction* in the *sim_true_counts* function, namely [0.6,0.75,0.9] to simulate different amounts of noise (lower values correspond to higher noise).

To generate the datasets in the latent simplex case, we first sampled three clusters and took their centroids as archetypes. The single cells were then simulated by sampling a matrix of archetype weights from a Dirichlet distribution **A** and multiplying it with the observation centroid for each modality **C**_*g*_. In the non-linear case, we first learn a latent space with a variational autoencoder **Z**, compute the centroids in this latent space, and then feed **AC** to the decoder (note that this time centroid are modality agnostic). We tested how well the methods reconstructed the archetype distribution **A**. If we call **Ã** the inferred the score we computed is:

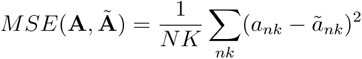

We also computed the Adjusted Rand Index (ARI) between the inferred and true highest archetype defined as 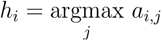.

For the trajectory cohorts, we were interested in comparing the archetypes to the terminal points of the trajectory. In this case, we define the terminal points as those having the lowest and highest pseudotime values. We computed a set of trajectory endpoints **t**_*k*_ by aggregating the expression of the bottom 15% percentile and the top 75% percentile of pseudotime for each terminal branch. We did the same to get and aggregate the 75% percentile of cells with the highest weight for each archetype to 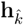. We matched each archetype index 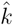 to the differentiation branch *k* with the lowest Euclidean distance and then computed:

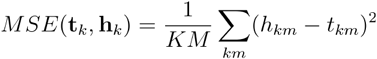

Where *M*is the number of features. We computed this score for both the RNA and the ATAC reconstruction.

### Real Data analysis: G&T

For the methylation and expression CD34+ dataset, we first filtered the CpG data by keeping only those with sites with less than 65% missing cells. We then filled the NA with 0 (unmethylated CpG). For the RNA we used as input the batch-correction latent representation of Scanorama ^32^ already computed by the authors in the original work. We then run our model with a Gaussian likelihood for the RNA and a Bernoulli likelihood for the methylation. We set a batch size of 300, a learning rate of 0.0001 with an exponential decaying schedule with a rate of 0.1, and run the inference for 1000 epochs using the Adam^33^ optimizer. We run the model for a number of archetypes ranging from 2 to 12 and choose the best value of 4 based on plateaus in the ELBO plot (Supplementary Figure 3).

Scores for the different progenitor cells were computed using the function *score_genes* of Scanpy^34^ from the gene sets in ^19^.

To compare the representation power of the different methods we set the number of latent dimensions in both the VAE and MOFA to 4 and correlate the gene scores to the latent coordinates. For the K-mean clustering we again chose 4 as the number of clusters, but this time we learned a MOFA model with 30 factors to simulate a more realistic scenario. GSEA^35^ was computed for archetypes 1 and 3 on the cell progenitor gene sets using the Python packages^36^.

### Real Data analysis:10x Multiome

The input matrices for RNA and ATAC where generated by taking the highly variable genes and 10000 peaks and then log transform and scale them after a library size normalization using Scanpy and SnapATAC^37^. We then run the model with a Gaussian likelihood for 3000 steps, exponential decay of 0.1. The best number of archetypes 6 was selected by again running the model in a range of [2,…,12] and looking at the negative ELBO decrease.

We confirmed the relation between archetype weights and cell fate commitment by first running chromVAR^21^ to obtain transcription factor deviation scores and then correlating marker TFs with archetype weights. We used the model learned from this dataset to generate some synthetic data. We sampled archetype weights for each cell from a Dirichlet with concentrations [1e-16,1e-16, 2,1, 2] that were then fed to the decoder.

## Data Availability

All the data used in this paper is publically available the CD34+ methylation and RNA datasets are stored on GEO and has accession number GSE158057 while the CD34+ 10x multiome can be downloaded from the Human Cell Atlas portal (https://explore.data.humancellatlas.org/projects/091cf39b-01bc-42e5-9437-f419a66c8a45)

## Code Availability

Our MIDAA tool is available as a Python package on PyPI, the source code is hosted on GitHub (https://github.com/sottorivalab/midaa). Scripts to reproduce the analysis are hosted at (https://github.com/sottorivalab/midaa_reproducibility)

## References

1. Simon, H. A. The Architecture of Complexity. Proc. Am. Philos. Soc. 106, 467–482 (1962).

2. Barabási, A.-L. & Oltvai, Z. N. Network biology: understanding the cell’s functional organization. Nat. Rev. Genet. 5, 101–113 (2004).

3. Karczewski, K. J. & Snyder, M. P. Integrative omics for health and disease. Nat. Rev. Genet. 19, 299–310 (2018).

4. Tarazona, S. et al. Harmonization of quality metrics and power calculation in multi-omic studies. Nat. Commun. 11, 3092 (2020).

5. Meng, C. et al. Dimension reduction techniques for the integrative analysis of multi-omics data. Brief. Bioinform. 17, 628–641 (2016).

6. Argelaguet, R. et al. MOFA+: a statistical framework for comprehensive integration of multi-modal single-cell data. Genome Biol. 21, 111 (2020).

7. Argelaguet, R. et al. Multi-Omics Factor Analysis—a framework for unsupervised integration of multi-omics data sets. Mol. Syst. Biol. 14, e8124 (2018).

8. Velten, B. et al. Identifying temporal and spatial patterns of variation from multimodal data using MEFISTO. Nat. Methods 19, 179–186 (2022).

9. Feist, A. M., Herrgård, M. J., Thiele, I., Reed, J. L. & Palsson, B. Ø. Reconstruction of biochemical networks in microorganisms. Nat. Rev. Microbiol. 7, 129–143 (2009).

10. Zuin, J. et al. Nonlinear control of transcription through enhancer-promoter interactions. Nature 604, 571–577 (2022).

11. Ibarra, I. L. et al. Mechanistic insights into transcription factor cooperativity and its impact on protein-phenotype interactions. Nat. Commun. 11, 124 (2020).

12. Igler, C., Rolff, J. & Regoes, R. Multi-step vs. single-step resistance evolution under different drugs, pharmacokinetics, and treatment regimens. Elife 10, (2021).

13. Lopez, R., Regier, J., Cole, M. B., Jordan, M. I. & Yosef, N. Deep generative modeling for single-cell transcriptomics. Nat. Methods 15, 1053–1058 (2018).

14. Ashuach, T. et al. MultiVI: deep generative model for the integration of multimodal data. Nat. Methods 20, 1222–1231 (2023).

15. Montavon, G., Samek, W. & Müller, K.-R. Methods for interpreting and understanding deep neural networks. Digit. Signal Process. 73, 1–15 (2018).

16. Cutler, A. & Breiman, L. Archetypal Analysis. Technometrics 36, 338–347 (1994).

17. Keller, S. M., Samarin, M., Wieser, M. & Roth, V. Deep Archetypal Analysis. in Pattern Recognition 171–185 (Springer International Publishing, 2019).

18. Paszke, A. et al. PyTorch: An imperative style, high-performance deep learning library. Adv. Neural Inf. Process. Syst. abs/1912.01703, (2019).

19. Nam, A. S. et al. Single-cell multi-omics of human clonal hematopoiesis reveals that DNMT3A R882 mutations perturb early progenitor states through selective hypomethylation. Nat. Genet. 54, 1514–1526 (2022).

20. Setty, M. et al. Characterization of cell fate probabilities in single-cell data with Palantir. Nat. Biotechnol. 37, 451–460 (2019).

21. Schep, A. N., Wu, B., Buenrostro, J. D. & Greenleaf, W. J. chromVAR: inferring transcription-factor-associated accessibility from single-cell epigenomic data. Nat. Methods 14, 975–978 (2017).

22. Mørup, M. & Hansen, L. K. Archetypal analysis for machine learning and data mining. Neurocomputing 80, 54–63 (2012).

23. Wang, Y. & Zhao, H. Non-linear archetypal analysis of single-cell RNA-seq data by deep autoencoders. PLoS Comput. Biol. 18, e1010025 (2022).

24. Persad, S. et al. SEACells infers transcriptional and epigenomic cellular states from single-cell genomics data. Nat. Biotechnol. 41, 1746–1757 (2023).

25. Hart, Y. et al. Inferring biological tasks using Pareto analysis of high-dimensional data. Nat. Methods 12, 233–5, 3 p following 235 (2015).

26. Kingma, D. P. & Welling, M. Auto-Encoding Variational Bayes. arXiv [stat.ML] (2013).

27. Davidson, T. R., Falorsi, L., De Cao, N., Kipf, T. & Tomczak, J. M. Hyperspherical Variational Auto-Encoders. arXiv [stat.ML] (2018).

28. Li, H., Zhang, Z., Squires, M., Chen, X. & Zhang, X. scMultiSim: simulation of single cell multi-omics and spatial data guided by gene regulatory networks and cell-cell interactions. Res Sq (2023) doi:10.21203/rs.3.rs-3301625/v1.

29. Lock, E. F., Hoadley, K. A., Marron, J. S. & Nobel, A. B. JOINT AND INDIVIDUAL VARIATION EXPLAINED (JIVE) FOR INTEGRATED ANALYSIS OF MULTIPLE DATA TYPES. Ann. Appl. Stat. 7, 523–542 (2013).

30. Chalise, P. & Fridley, B. L. Integrative clustering of multi-level ‘omic data based on non-negative matrix factorization algorithm. PLoS One 12, e0176278 (2017).

31. Eugster, M. & Leisch, F. From Spider-man to Hero - archetypal analysis in R. 1 (2009).

32. Hie, B., Bryson, B. & Berger, B. Efficient integration of heterogeneous single-cell transcriptomes using Scanorama. Nat. Biotechnol. 37, 685–691 (2019).

33. Kingma, D. P. & Ba, J. Adam: A Method for Stochastic Optimization. arXiv [cs.LG] (2014).

34. Wolf, F. A., Angerer, P. & Theis, F. J. SCANPY: large-scale single-cell gene expression data analysis. Genome Biol. 19, 15 (2018).

35. Subramanian, A. et al. Gene set enrichment analysis: a knowledge-based approach for interpreting genome-wide expression profiles. Proc. Natl. Acad. Sci. U. S. A. 102, 15545–15550 (2005).

36. Fang, Z., Liu, X. & Peltz, G. GSEApy: a comprehensive package for performing gene set enrichment analysis in Python. Bioinformatics 39, (2023).

37. Fang, R. et al. Comprehensive analysis of single cell ATAC-seq data with SnapATAC. Nat. Commun. 12, 1337 (2021).

