## Supplementary Information for "Deep Archetypal Analysis for interpretable multi-omic data integration based on biological principles"

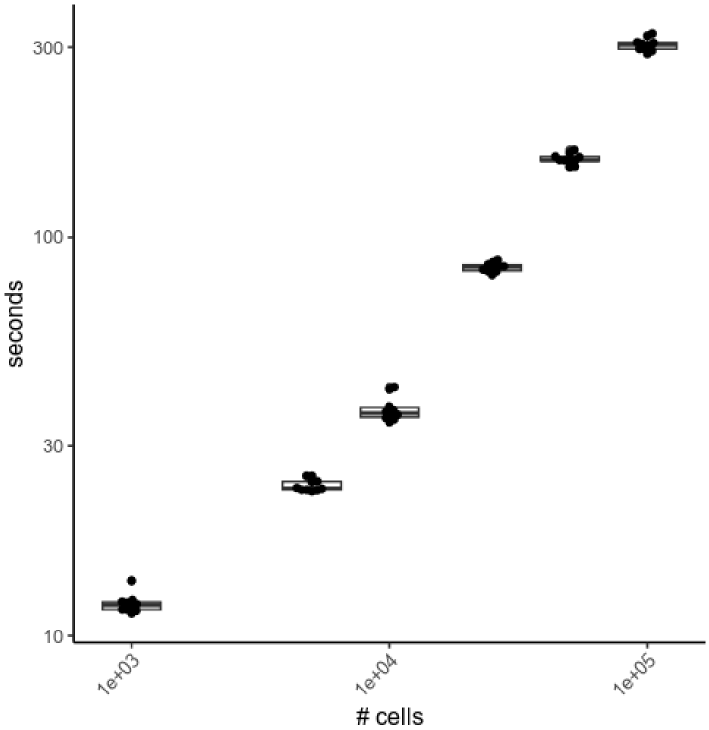


**Supplementary Figure 1.** Running time for MIDAA. We run 10 instances of the model for each number of cells for 500 epochs with a batch size of 4096. Tests were run on a CUDA backend on an NVIDIA V100 SMX2 with 32 GB of dedicated RAM.

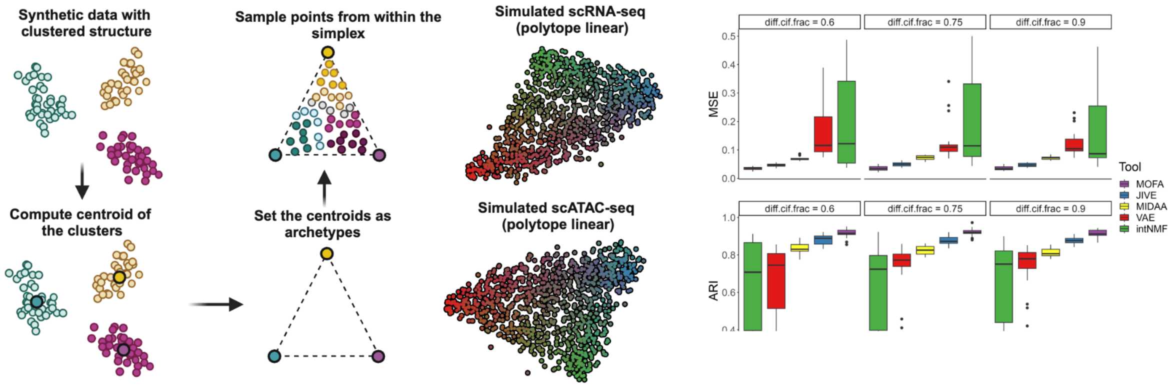


**Supplementary Figure 2.** A simple test case consists of reconstructing a simplex structure in a linear latent space. The UMAP clearly shows the latent polytope structure. On the right: top panel is the Mean Squared Error (MSE) between the true archetype weight matrix and the inferred one. The bottom panel is the Adjusted Rad Index (ARI) between the true archetype assignments and the inferred ones.

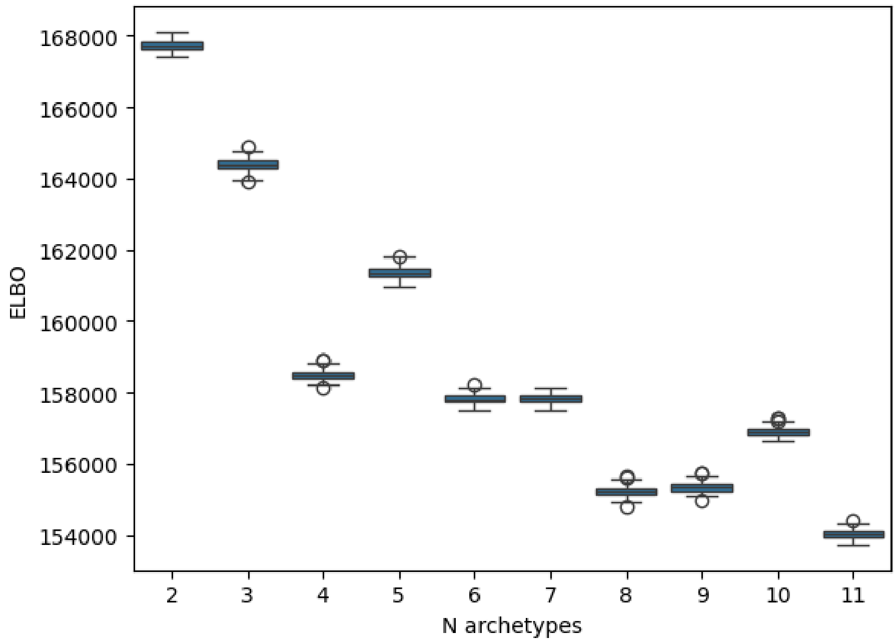


**Supplementary Figure 3. ELBO plateau for the G&T dataset.** We plotted the ELBO value for the last 50 steps of the inference. We choose 4 as the optimal number of archetypes as it corresponds to the first plateau in the loss.

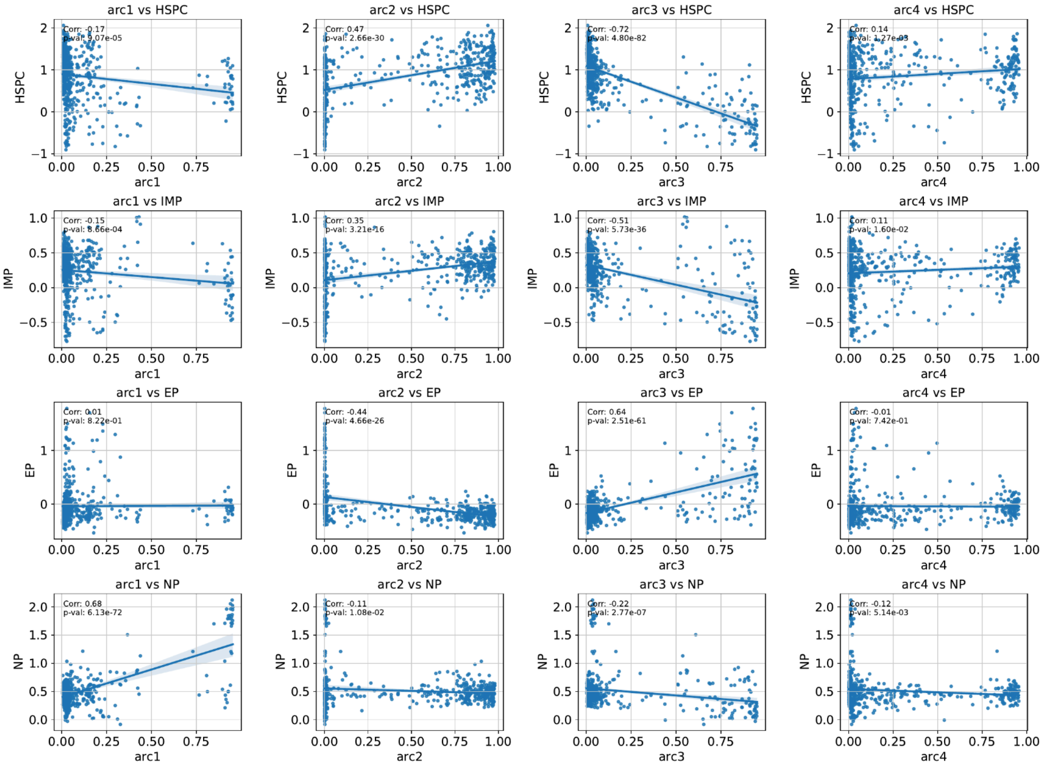


**Supplementary Figure 4. Correlation between progenitor lineage score and archetype weights.** Interestingly we see how archetype one is correlated strongly with NP cells, while archetype 2 correlates with the EP lineage. Archetype 2 seems to be correlated with stem lineages, which are expectedly anticorrelated in archetypes 1 and 3. Archetype 4 seems to have captured signal not related to the lineage commitment process.

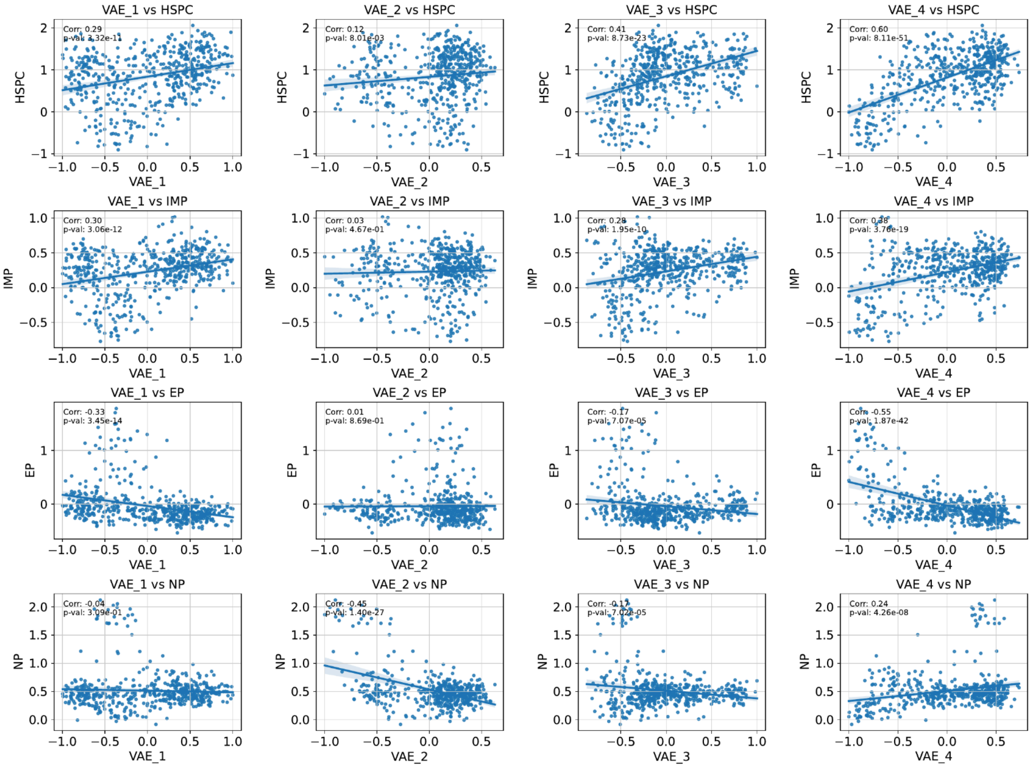


**Supplementary Figure 5. Correlation between progenitor lineage score and archetype weights.** While the coordinates of the VAE latent space show some degree of correlation especially with the stem commitment program (which is the principal one in the dataset) they fail to recapitulate cell commitment towards NP and EP cells.

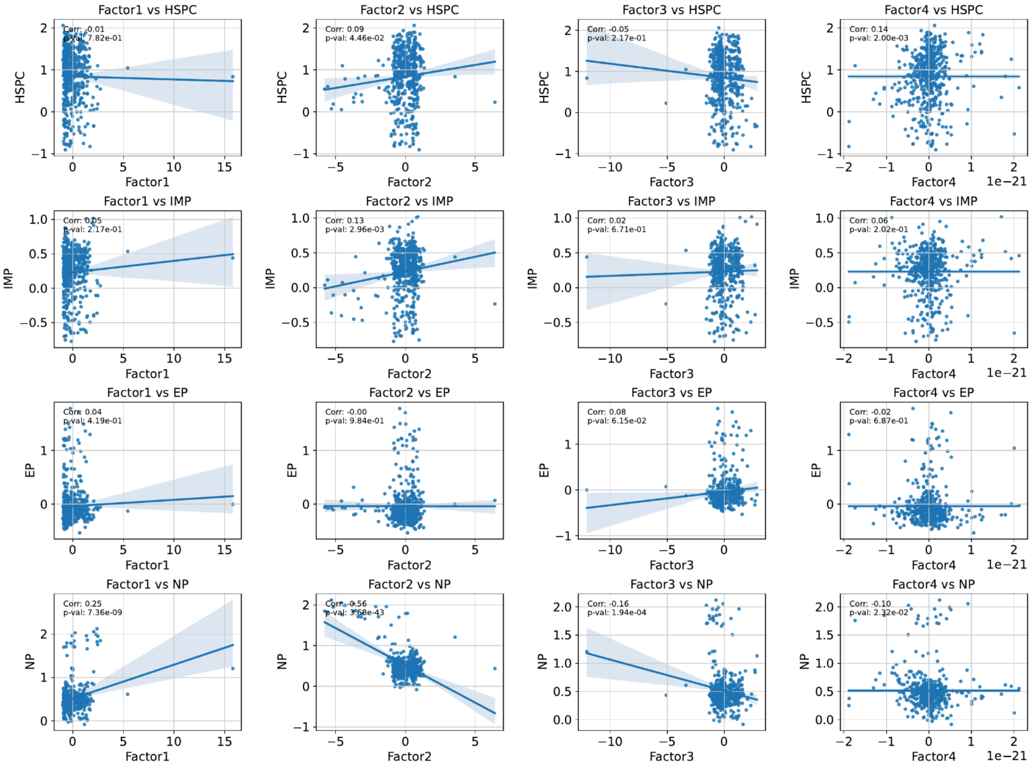

**Supplementary Figure 6. Correlation between progenitor lineage score and archetype weights.** Here the first 4 MOFA factors are correlated with some high variance points, this shows how MOFA needs more coordinates to produce an efficient representation of the dataset in latent space. This is also related to the fact that factors in MOFA are ordered according to the variance explained.

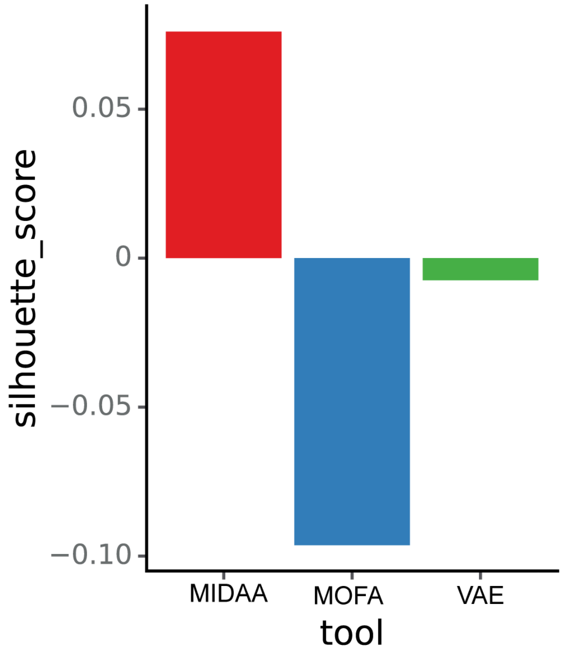


**Supplementary Figure 7. Silhouette scores for the clustering strategies in Figure 2 C.** We compute the silhouette scores using the scores for the terminal progenitor states of the dataset, namely EP, NP, and HSPC. MIDAA's increase in Silhouette score quantitatively confirms what can be seen in Figure 2C.

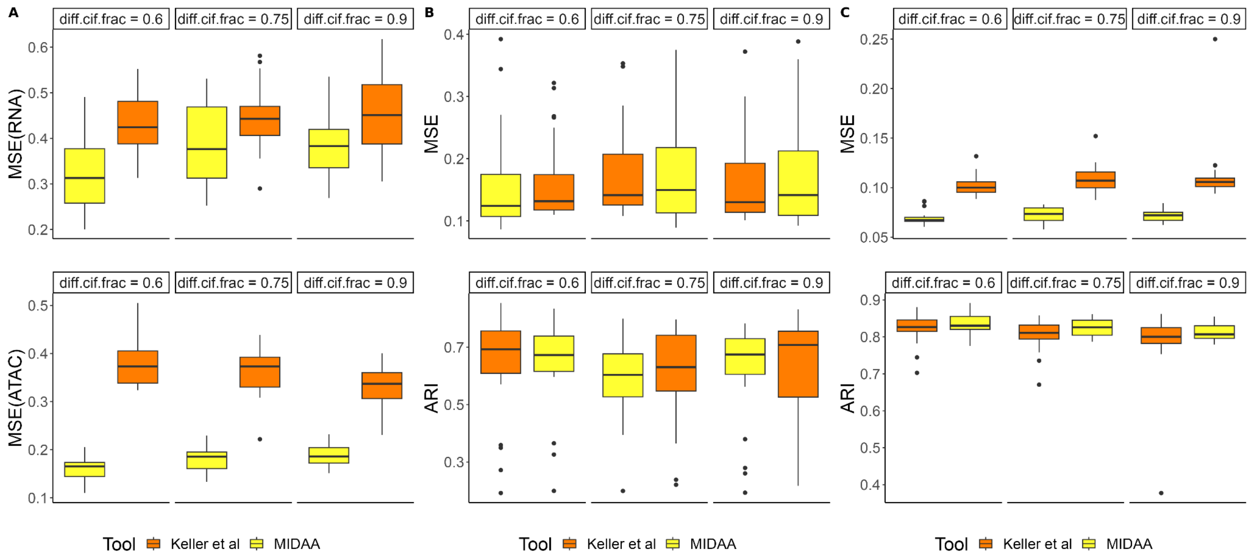


**Supplementary Figure 8. Difference in performance of the latent space of MIDAA and of Keller et al. 2019[17].** Panels A, B, and C correspond to the tests in Figure 1 D, G, and Supplementary Figure 3 respectively. We see how the MIDAA model on average outperforms the formulation of Keller et al. 2019[17] in this set of tests.
